# Dynamic functional brain networks underlying the temporal inertia of negative emotions

**DOI:** 10.1101/2021.03.26.437275

**Authors:** Julian Gaviria, Gwladys Rey, Thomas Bolton, Dimitri Van De Ville, Patrik Vuilleumier

## Abstract

Affective inertia represents the lasting impact of transient emotions at one time point on affective state at a subsequent time point. Here we describe the neural underpinnings of inertia following negative emotions elicited by sad events in movies. Using a co-activation pattern analysis of dynamic functional connectivity, we examined the temporal expression and reciprocal interactions among brain-wide networks during movies and subsequent resting periods. Our findings revealed distinctive spatiotemporal expression of visual (VIS), default mode (DMN), central executive (CEN), and frontoparietal control (FPCN) networks both in negative movies and in rest periods following these movies. We also identified different reciprocal relationships among these networks, in transitions from movie to rest. While FPCN and DMN expression increased during and after negative movies, respectively, FPCN occurrences during the movie predicted lower DMN and higher CEN expression during subsequent rest after neutral movies, but this relationship was reversed after the elicitation of negative emotions. Changes in FPCN and DMN activity correlated with more negative subjective affect. These findings provide new insights into the transient interactions of intrinsic brain networks underpinning the inertia of negative emotions. More specifically, they describe a major role of FPCN in emotion elicitation processes, with prolonged impact on DMN activity in subsequent rest, presumably involved in emotion regulation and restoration of homeostatic balance after negative events.

**Highlights:**

- Modulations of dynamic functional brain connectivity are associated to the temporal inertia of negative emotions.
- Functional co-activations patterns (CAPs) during emotional episodes predict changes in spontaneous brain dynamics during subsequent resting state.
- Classical “task-rest” anticorrelations in network activity are reversed by negative emotions.

## Introduction

Abundant work in neuroscience shows that functional brain activity and connectivity is dynamically organized across several distributed networks with simultaneous ongoing fluctuations and selective reciprocal interactions (e.g., correlated or anticorrelated) according to current behavioral demands (Fox et al., 2005). These spatiotemporal fluctuations in intrinsic functional networks (IFNs) can be observed during particular tasks as well as during rest. Among IFNs, the default mode network (DMN) encompasses a set of midline cortical areas, including precuneus and medial prefrontal cortices (MPFC) that are usually active during mind-wandering in “task-off” conditions at rest (Christoff et al., 2016). However, this network might also play a more specific and active role in integrative functions related to higher-level aspects of self-awareness and introspection (Fair et al., 2008). The DMN activity is associated with self-reflective and memory-related task conditions (Spreng et al., 2009) and typically anticorrelates with activity of the frontoparietal control network (FPCN), which is recruited by externally directed attention (Dixon et al., 2018a). DMN activity also fluctuates between periods of positive and negative correlation with the central executive network (CEN) and saliency networks (SN), indicating switches between different cognitive processing modes (Dixon et al., 2017). In addition, there is evidence that the DMN expression is modulated by emotional information and current mood (Gaviria et al., 2021; Nummenmaa et al., 2012; Satpute and Lindquist, 2019). However, its exact role in affective processes is unclear.

In the present study, we investigated how IFNs dynamically reconfigure and reciprocally interact with each other in response to negative emotional episodes, both during emotional stimulation itself and during the recovery period following such stimulation. Dynamic reciprocal interactions among IFNs dovetails with a general theoretical framework of affective phenomena (Scherer, 2009) according to which emotions emerge from a coordinated recruitment of component processes, each underpinned by distinct neural systems, whose synchronization is triggered by behaviorally relevant events and promotes adaptive changes in the organism (Leitão et al., 2020; Meuleman and Rudrauf, 2018). Evidence for affective influences on DMN activity and its interaction with other networks comes from several observations (Buckner and DiNicola, 2019; Kaiser et al., 2015), but their functional significance remains unresolved. A down-regulation of within-DMN connectivity was reported in healthy participants during sad mood induction through self-generated memories (Harrison et al., 2008) or movies (Borchardt et al., 2018), while disturbances of DMN connectivity are observed in clinical mood disorders such as depression and bipolar disease (Rey et al., 2014; Zovetti et al., 2020). Dynamic shifts in the balance between DMN and SN are also thought to mediate adaptive responses to acute stressors, promoting higher vigilance and fear (Zhang et al., 2019). Furthermore, changes in the activation and spatial configuration of DMN at rest can be modified by preceding cognitive (Barnes et al., 2009) or affective (Eryilmaz et al., 2014, 2011; Gaviria et al., 2021) task conditions. In particular, exposure to emotional stimuli or rewards has been found to induce long-lasting changes in brain activity and connectivity, overlapping with DMN, SN, and FPCN, which persist even after termination of the eliciting event itself, e.g., during subsequent resting state (Borchardt et al., 2018; Eryilmaz et al., 2011; Harrison et al., 2008; van Marle et al., 2010). Such carry-over effects of emotions on neural activity at rest might reflect spontaneous self-regulation mechanisms restoring a homeostatic balance in brain state (Eryilmaz et al., 2011; Gaviria et al., 2021), and is enhanced by individual anxiety levels (Pichon et al., 2015). This is consistent with behavioral evidence for a temporal “inertia” of affect following transient emotions, exacerbated by maladaptive emotion regulation (Kuppens and Verduyn, 2017), and contributing to ruminative processes triggered by negative events in psychopathological conditions (Apazoglou et al., 2019). In accordance with this idea, cognitive reappraisal of negative audiovisual stimuli (compared to freely watching) may produce not only a reduced neural response to these stimuli in limbic areas (e.g., amygdala), but also an increased activation of cortical networks overlapping with DMN and FPCN during both stimulus presentation and subsequent rest (Lamke et al., 2014).

However, previous studies on emotion-responsive networks and emotion carry-over generally focused on *static* connectivity patterns, temporally summarized at the whole-brain level, either during emotional stimulation or during resting conditions following emotional stimulation (compared to neutral). They did not characterize the *dynamics* of changes in specific IFNs during these different periods; nor did they examine how these changes may emerge from interactions *between* different IFNs simultaneously modulated by emotional state. Hence, these studies do not offer any mechanistic insights into how IFNs dynamically unfold during and after emotions, and how their acute engagement by an emotion-eliciting event may influence the subsequent return to a normal resting baseline after termination of this event (Hermans et al., 2014).

To directly address these issues, we designed a novel emotion elicitation paradigm where participants freely viewed sad (vs. neutral) movies followed by a resting state period. To uncover IFNs differentially engaged during emotional movies and their subsequent carry-over at rest, as well as their reciprocal relationships, we leveraged an innovative methodology allowing us to track dynamic functional connectivity (dFC) with co-activation pattern (CAP) analysis (Liu et al., 2018b; Liu and Duyn, 2013). By delineating brain-wide CAPs modulated by emotion in movie and rest periods, we asked the following questions: (1) what is the carry-over impact of negative events on the dynamics and reciprocal interactions of brain networks and how do they unfold in time beyond these events, i.e., in the aftermath of emotion during subsequent rest (see Fig 1A); (2) whether changes in brain states subsequent to negative emotions, particularly in DMN and interconnected networks, are associated with changes in subjective affective state; and (3) how is the carry-over effect emotion on resting IFNs related to the occurrence of specific activity patterns during the preceding emotional events. By characterizing the spatiotemporal features of IFNs during and after negative affect induction, our study provides new insights on the brain dynamics of emotional responses and may help identify novel neurobiological markers for negative affect experience and regulation, possibly altered in clinical populations with anxiety and mood disorders.

**Figure 1.**
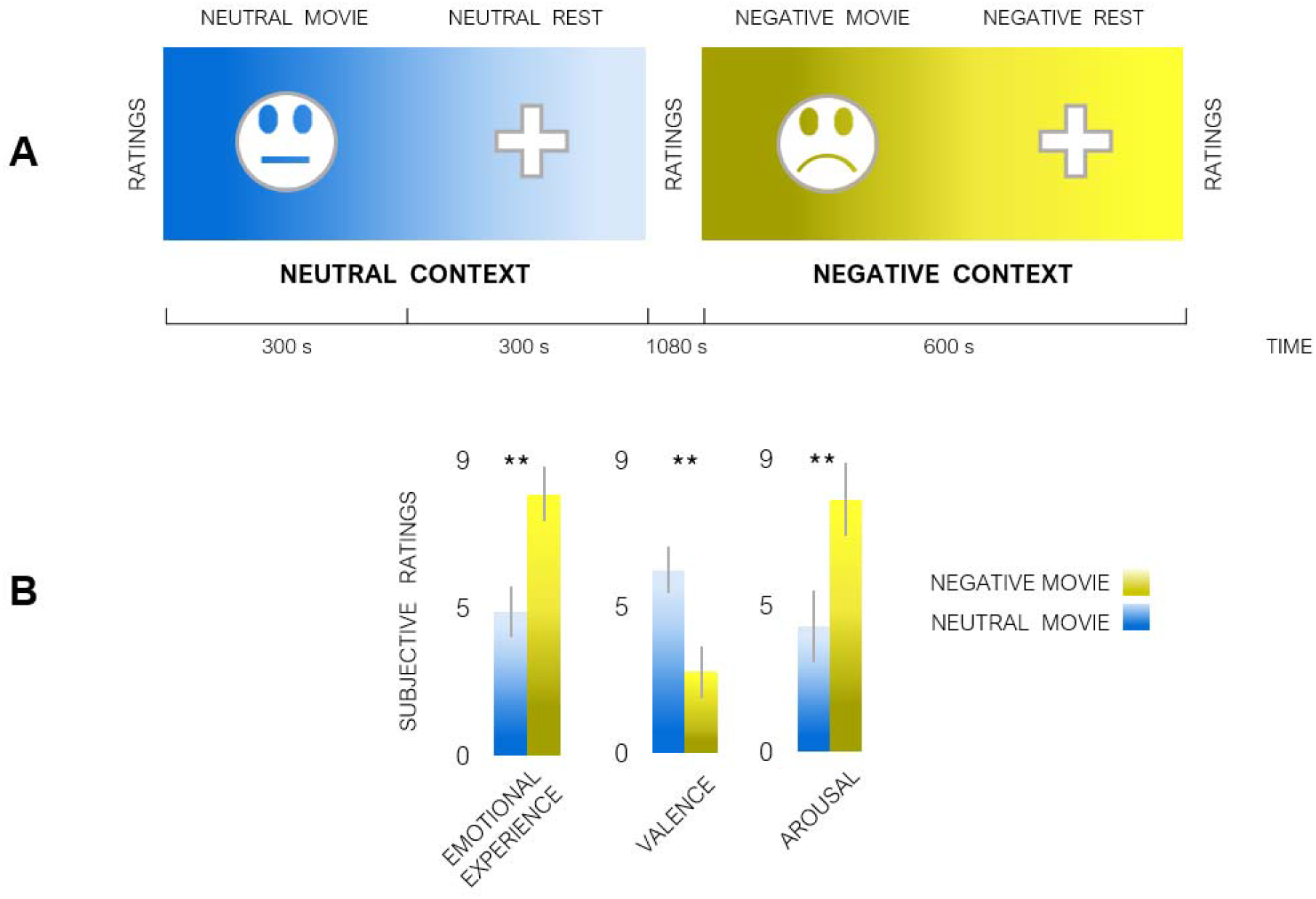
Behavioral paradigm. **A)** Paradigm design illustrating the sequence of experimental events. Negatively-valenced or neutral movies were followed by a resting period. The presentation of the two affective contexts (i.e., “neutral” and “negative”) was counterbalanced across participants. **B)** Subjective affective ratings of movies obtained after each experimental context. Valence was rated from very negative on the lowest scale values (=0) to very positive on the highest scale value (=9). Whiskers stand for standard errors of means. P_FDR_ adjustment for multiple comparisons (** p<.01).

## Materials and Methods

### Participants

Twenty right-handed, French-speaking female volunteers (mean age= 23.2 ± 4.3) were contacted via posters and online advertisement. They all provided written informed consent according to the Geneva University Hospital Ethics Committee. Only female participants were recruited because pilot testing suggested stronger emotional induction in women, compared to male, particularly with the movie clips used here (see Supplementary methods). Controlled inclusion criteria were: no history of neurological and psychiatric diseases; known menstrual phase and no contraceptive method to rule out hormonal effects on emotional processing (Protopopescu et al., 2005) or functional connectivity (Petersen et al., 2014; Pletzer et al., 2016); no major head movement during the scanning sessions (<1.5 mm in all axes). Scanning was scheduled for each participant in the early stage of the menstrual follicular phase, when the levels of estradiol and progesterone hormones are moderate (Andreano et al., 2018; Endicott, 1993). Nicotine and caffeine consumption was prohibited 10 hours before scanning.

### Behavioral paradigm

Initially, the participants filled the French version of the PANAS questionnaires. This was followed by an fMRI experiment with a repeated measure design. The scanning session comprised two experimental “contexts”. One context included an audio-visual clip (5 min) with sad emotional content (“negative movie” condition), followed by a resting period (5 min) (“negative rest” condition). As a control for affective valence, the second experimental context comprised non-emotional video clip (5 min) (“neutral movie”), followed by another resting period (“neutral” rest condition) (5min). Participants were instructed to keep their eyes opened and to stay awake across the resting periods. This was controlled by an eye-tracking device both online and offline. Importantly, the presentation order of the two contexts was counterbalanced across participants (see Figure 1A, for schematic overview of the paradigm design). At the end of each experimental scanning session (i.e., “neutral” and “negative”), further PANAS scores and emotion questionnaires about the movies were also filled. Approximately 20 minutes elapsed between the end of one experimental context and the starting of the next one. Additional information on movies and subjective ratings is provided in the SI material.

### MRI data acquisition

Anatomical and functional whole-brain MRI data were acquired with a 3T scanner (Siemens TIM Trio) at the Brain & Behavior Laboratory, University of Geneva. A multiband-sequence was implemented, with a voxel size of 2mm isometric and a TR of 1.3s (see SI for further description of both data acquisition and preprocessing).

### Brain data analysis

The present study focused on the dynamic expression and interaction of IFNs over time both during stimulation with negative movies and at rest following stimulation (i.e., emotional inertia). To this aim, our approach comprised two successive stages with different goals: (1) Preliminary GLM-based identification of brain regions of interest (ROIs) with functionally relevant activity during our paradigm to be used as seeds; (2) In-depth CAP analysis of the time-varying dynamic functional connectivity of these seeds to characterize brain networks differentially engaged by experimental conditions, as well as their interactions at two different time points (i.e., movie and rest). This approach entails a two-step data-driven approach that integrates two different and independent levels of interrogation of brain activity in our task-rest paradigm (Shine and Poldrack, 2018). Namely, an assessment of functional activation level (GLM) along a magnitude scale, and dynamic functional connectivity level (CAPs) along a temporal domain.

### Functional seed definition by univariate analysis of brain activity

We first performed a GLM analysis to define ROIs as seeds for the main CAP analysis. Changes in global neural activity were assessed between sad and neutral conditions during movies (“negative movie” vs. “neutral movie” conditions) and during rest (“negative rest” vs. “neutral rest” conditions) using a standard procedure for block design in SPM12 (Frackowiak et al., 1997). In addition to removing signals at very low frequencies, temporal autocorrelation across the functional data was accounted for by using the “FAST” algorithm in SPM12 (Corbin et al., 2018; Olszowy et al., 2019). Four GLM boxcar regressors represented each experimental condition (i.e., neutral movie, neutral rest, negative movie, and negative rest respectively). These regressors were convolved with a standard hemodynamic response function (HRF) according to a blocked design, which was then submitted to a univariate regression analysis. Realignment parameters were added to the design matrices of both models, to account for any residual movement confounds. In all cases, the design matrix included low-frequency drifts with a cutoff frequency at 1/128 Hz (other cutoff frequencies did not change the GLM results). Flexible factorial analyses of variance (ANOVAs) were then performed on the main contrasts of interest (i.e., “negative movie > neutral movie”, and “negative rest > neutral rest”), to define two different seed ROIs for our main connectivity analyses. One with stimulus-related activity and another with rest-related activity. All statistical analyses were carried out at the whole-brain level, with a threshold at P_FWE_<.05, corrected at whole brain level (unless specified otherwise).

### Dynamic functional connectivity analysis (CAPs identification)

To examine changes in IFNs as a function of experimental conditions (neutral movie, neutral rest, negative movie, and negative rest), we implemented a co-activation pattern (CAP) methodology (Chang et al., 2016; Liu and Duyn, 2013; Preti et al., 2017). It has been previously shown (Liu et al., 2013; Liu and Duyn, 2013) how transient associations between fMRI signals captured by specific co-activation patterns (CAPs), correspond to the intrinsic neural architecture configured for supporting specialized functions (IFNs). Remarkably, the CAP approach provides a measure of dynamic functional connectivity with a relevant seed by decomposing fluctuations of the fMRI BOLD signals into multiple spatial patterns that reflect the instantaneous organization and reorganization of networks co-activated with the seed over time (Liu et al., 2018b; Preti et al., 2017). The two emotion-responsive ROIs identified by our GLM analysis (insula and precuneus from “negative movie” and “negative rest” conditions, respectively) were used as seeds for the CAPs generation. By using seeds defined by an independent GLM-based analysis, we could ensure to “anchor” the observed CAPs to functionally relevant networks in the current experimental paradigm, rather than highlight other less specific IFNs. Unlike static connectivity analysis based on correlations by averaging over long periods, the dFC approach with CAPs accounts for the BOLD temporal variability by representing instantaneous brain configurations at single time points (fMRI volumes) and then sorting them in a few dominant patterns through clustering analysis (Liu et al., 2018b). CAPs were computed with the “tbCAPs” toolbox (Bolton et al., 2020), following a stepwise pipeline illustrated in Figure S1A more detailed description of our methodological procedure is provided elsewhere (Bolton et al., 2020). Importantly, this approach yields a temporal metric to quantify dFC variability over time by computing the occurrences of each CAP in each condition. Occurrences are defined as the sum of frames (time-points) assigned to a given CAP among all the retained frames, across the entire scanning duration.

Following the generation of CAPs for each seed, we compared their expression across the affective contexts with generalized linear mixed models [gLMM (Brooks et al., 2017)] using the R software. We performed a 2×2 gLMM-based factorial analysis comprising two fixed factors denoted by the following R-based formula syntax:

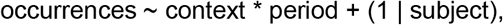

where the fixed factor “context” comprised two levels corresponding to the affective valence of movies (i.e., negative or neutral) and the second factor “period” represented the type of condition (movie or rest). Individual participants (subject) were modeled as a random factor, and “occurrences” was the dependent variable, computed for each experimental condition and each CAP. This 2×2 analysis allowed us to evaluate separately the main effects of stimulus exposure (movie vs. rest) and emotional valence (negative vs. neutral), as well as their interactions, for each of the CAPs. Subsequently, only the significant CAPs showing either main effects or interactions of the factors “context” (neutral, negative) and period (movie, rest), were considered as experimentally relevant and selected for more detailed evaluation (see next section).

### Association of affective scores with CAPs

We also tested whether differences in the temporal metrics of these relevant CAPs were associated with behavioral measures of subjective emotional state (PANAS). In doing so, we verified that all the PANAS data was normally distributed, except for the positive affective scores (PA) following the neutral context [Shapiro-Wilk test (p=.03)]. This was also the case for all metric from the different CAPs in both experimental contexts, except for the prec-VIS in “neutral rest” (p=.04); and prec-CEN in “neutral movie” (p=.02). In order to properly model these data, we implemented our correlation analyses with generalized estimated equations (GEEs), a statistical framework for flexibly modelling non-independent and correlated data (i.e., repeated measures) with both normal and non-normal distributions (Pekár and Brabec, 2018). In these analyses, the CAPs occurrences were introduced as multiple outcomes, and affective indices (PANAS scores) were the predictor variables. P values were adjusted with false discovery rate (FDR) for multiple testing (Benjamini and Hochberg, 1995), which is also optimal to control for dependency (Benjamini and Yekutieli, 2001).

### CAPs interaction analysis

#### Context-dependent within-CAPs interactions

To address the main goal of our study concerning functional relationships between networks in different affective states, we examined how changes in the expression of a given network correlated with changes in other networks across successive periods (e.g., movies vs. post-movie rest). First, we computed a set of within-CAP pair-wise Pearson coefficients (r) to assess the occurrence of each affectively-relevant CAP across different conditions. This analysis thus probed for context-dependent “movie-rest” transitions where occurrences during the “movie” period were correlated with occurrences during the subsequent “rest” period of the same CAP in the same affective valence [e.g., r(CAP(A)_neutral movie_, CAP(A)_neutral rest_) vs. r(CAP(A)_negative movie_, CAP(A)_negative rest_)]. As a control comparison, these within-CAP relationships were compared to movie-rest correlations between different contexts, when movies and rest conditions did not follow in direct succession [e.g., r(CAP(A)_negative movie_, CAP(A)_neutral rest_) vs. r(CAP(A)_neutral movie_, CAP(A)_negative rest_). Figure 4]. Because only reporting differences in significance for different conditions [e.g., one correlation significant (p<.05) for transitions in one context but not the other (p>.05)] would lead to statistical fallacy (Nieuwenhuis et al., 2011), we could thus directly assess differences in the magnitude of context/valence-specific correlations relative to non-specific control correlations by comparing the confidence intervals (CI) of their Pearson coefficients, which allows considering both the magnitude and precision of the estimated effects (Zou, 2007). All within-CAP comparisons were implemented with the R-based Cocor package (Diedenhofen and Musch, 2015). Dependency of within-CAP correlations were accounted following Zuo et al., (Zou, 2007), when their comparisons included overlapping variables [e.g., r(CAP-A_neutral movie_, CAP-A_neutral rest_) vs. r(CAP-A_neutral movie_, CAP-A_negative rest_). Figure 4].

#### Context-dependent between-CAPs interactions

A second set of analyses concerned the between-CAPs of interest associations, and allowed us to examine their reciprocal relationships as a function of emotional context. The correlation between occurrences of different CAPs was again computed context-wise, to probe how expression of one CAP during the movie watching period was associated to the expression of other CAPs in the just following resting state period [e.g., r(CAP(A)_neutral_, CAP(B)_post neutral_)]. The correlations for each pair of CAPs were then compared between the two affective contexts [e.g., r(CAP(A)_neutral_, CAP(B)_post neutral_) vs. r(CAP(A)_negative_, CAP(B)_post negative_). Figure 5B]. Dependency of non-overlapping variables was accounted for these between-CAPs comparisons as appropriate (Zou, 2007). Further description of statistical tests including R-based mathematical formulae for both within-CAPs and between-CAPs comparisons are described in “SI. Tests for comparing CAPs correlations” section.

## Results

### Behavioral indices of emotional induction by movies

We first verified that movies with negative content induced distinctive patterns in both behavior and brain measures, compared to neutral movies. Affective ratings of movie clips confirmed a reliable difference in emotional experience with more negative valence and higher arousal elicited by the negative compared to neutral movies (Figure 1B). PANAS scores assessing subjective affect also differed between the pre- and post-context measures as a function of emotional condition, with significant increases in the negative affect (NA) scores following negatively-valenced movies (Figure 3A). Further behavioral results concerning the movies are described in “SI. Psychological indices of emotion elicitation” section.

### Preliminary GLM-based definition of regions of interest

Prior to the CAP analysis, we defined functionally responsive seeds with stimulus-related and rest-related activity by computing linear contrasts between experimental blocks. Movies with negative compared to neutral valence produced selective increases in several limbic regions (P_FWE_<.05, whole brain-corrected at cluster level; cluster size estimated at P_uncorrected_<.001) predominantly in the left insula cortex (including both posterior and anterior parts, with peak MNI: −40, 4, 10; 121 voxels), together with the bilateral anterior cingulate cortex, bilateral middle frontal gyrus, and right temporal pole. As the insula has been consistently associated to emotional processing across many paradigms (Benelli et al., 2012; Critchley and Garfinkel, 2017; Liu et al., 2012; Paulus et al., 2005; Somerville et al., 2013), most often in relation to negative valence (Knutson et al., 2014), this region was selected as our first ROI (stimulus-related). Conversely, “negative rest” vs. “neutral rest” activated widespread areas associated with the DMN, predominantly in bilateral precuneus (peak MNI: 4, −54, 42; 541 voxels. P_FWE_<.05, whole brain-corrected at cluster level; cluster size estimated at P_uncorrected_<.001), and to a lesser degree in occipital cortex and left medial superior frontal gyrus. This accords with previous work reporting activation of the precuneus and other DMN regions at rest and their modulation by emotional or cognitive factors (Fox et al., 2018; Ho et al., 2015; Northoff, 2016; Spreng et al., 2010). We therefore selected the precuneus as the second seed (rest-related) for our subsequent connectivity analysis. Notably, our two seed ROIs largely overlapped with the insula (87% of voxels) and precuneus (78% of voxels) from a functional brain parcellation atlas (Schaefer et al., 2018), and converged with independent meta-analytic data showing consistent activation of these areas in emotion and rest conditions, respectively.

### CAPs expressed across stimulation periods and affective contexts

Two CAP analyses were conducted using the seed ROIs identified by our preliminary GLM analysis (see “Methods” section and Figure S1A), one reflecting functional coupling with the left insula and another reflecting coupling with the precuneus. In total 14 CAPs were identified, based on a data-driven “consensus” procedure (Monti et al., 2003) indicating K=7 for each seed as the optimal number of distinct brain maps in terms of replicability for our dataset (see pipeline in Figure S1). The spatial configuration of these CAPs is depicted in “SI. Figure S2.2” and globally accords with previous work on IFNs (Christoff et al., 2016; Cole et al., 2014; Goldman-Rakic, 1995; Kragel et al., 2019; Uddin et al., 2019).

### Changes in temporal dynamics of CAPs across conditions

Critically, we then compared the temporal occurrences of all the 14 CAPs between experimental conditions. Using a 2 (affective context) x 2 (stimulation period) factorial analysis of CAP occurrences, we found that only 4 out of these 14 networks were differentially expressed across the affective contexts (i.e., negative vs. neutral. Table 1), including the insula-connected CAP5 (insuFPCN) and the precuneus-connected CAP1, CAP3, and CAP5 [precVIS, precDMN, and precCEN, respectively (Figure.2)]. Peak coordinates for these four networks are listed in “SI. Table S1”. Below we describe each of these CAPs in turn.

**Table 1.**
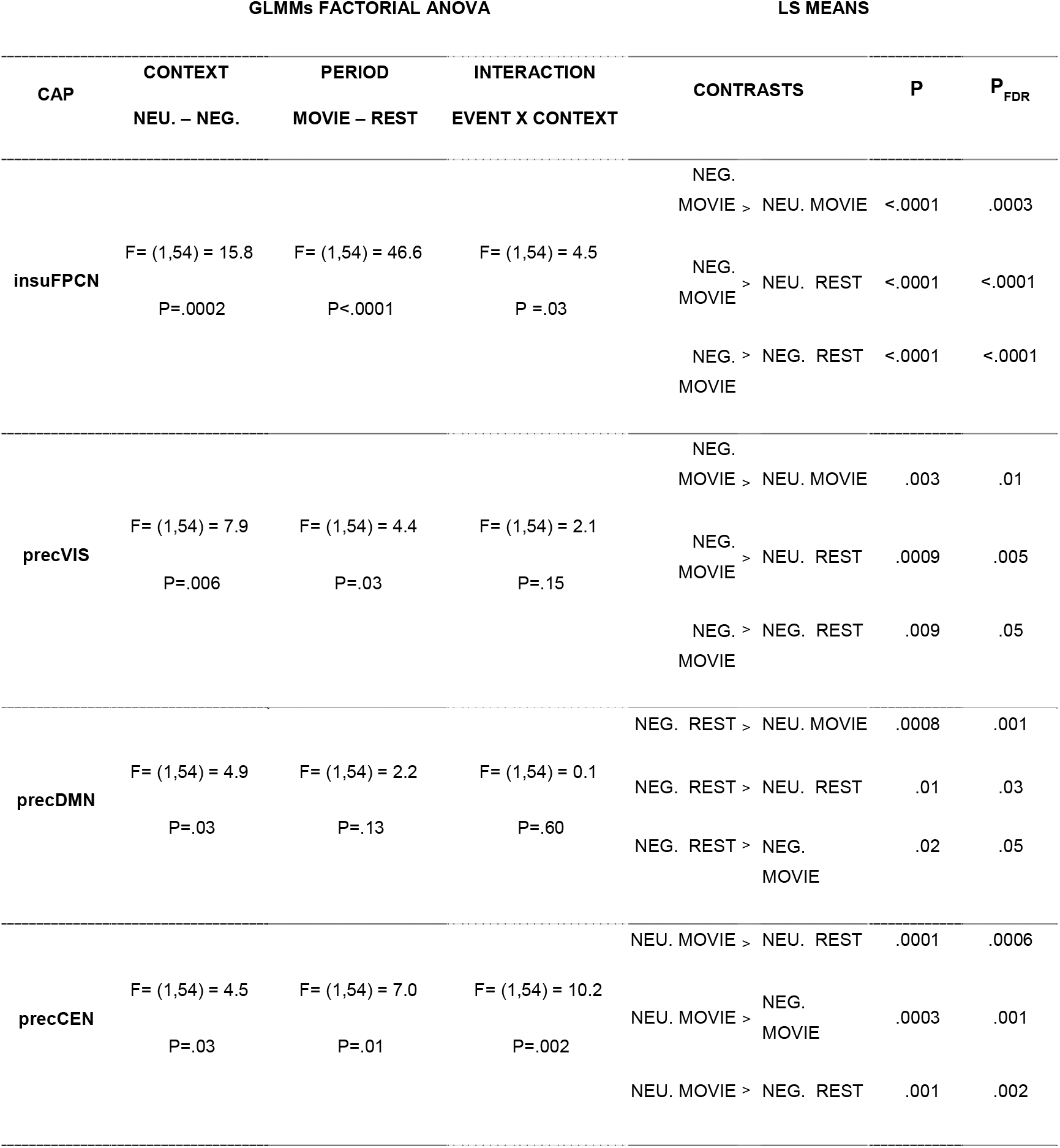
Results of GLMMs analyses on temporal occurrences of relevant / emotionsensitive CAPs. Statistical results of a 2×2 factorial assessment of the “context” (neutral vs. negative), and “period” (movies vs. rest) effects on CAPs occurrences. Significant post hoc pairwise contrasts driving main effects and interactions are listed in the right-hand columns., this factorial analysis revealed that 4 out of the 14 brain networks identified in our CAP analysis (see full description in Fig. S2) exhibited a distinctive activity profile according to our experimental conditions. These four relevant CAPs were therefore selected for all main analyses in our study. P_FDR_ (false discovery rate) adjustment for multiple comparisons (* p<.05; ** p<.01; ***p<.001).

**Figure 2.**
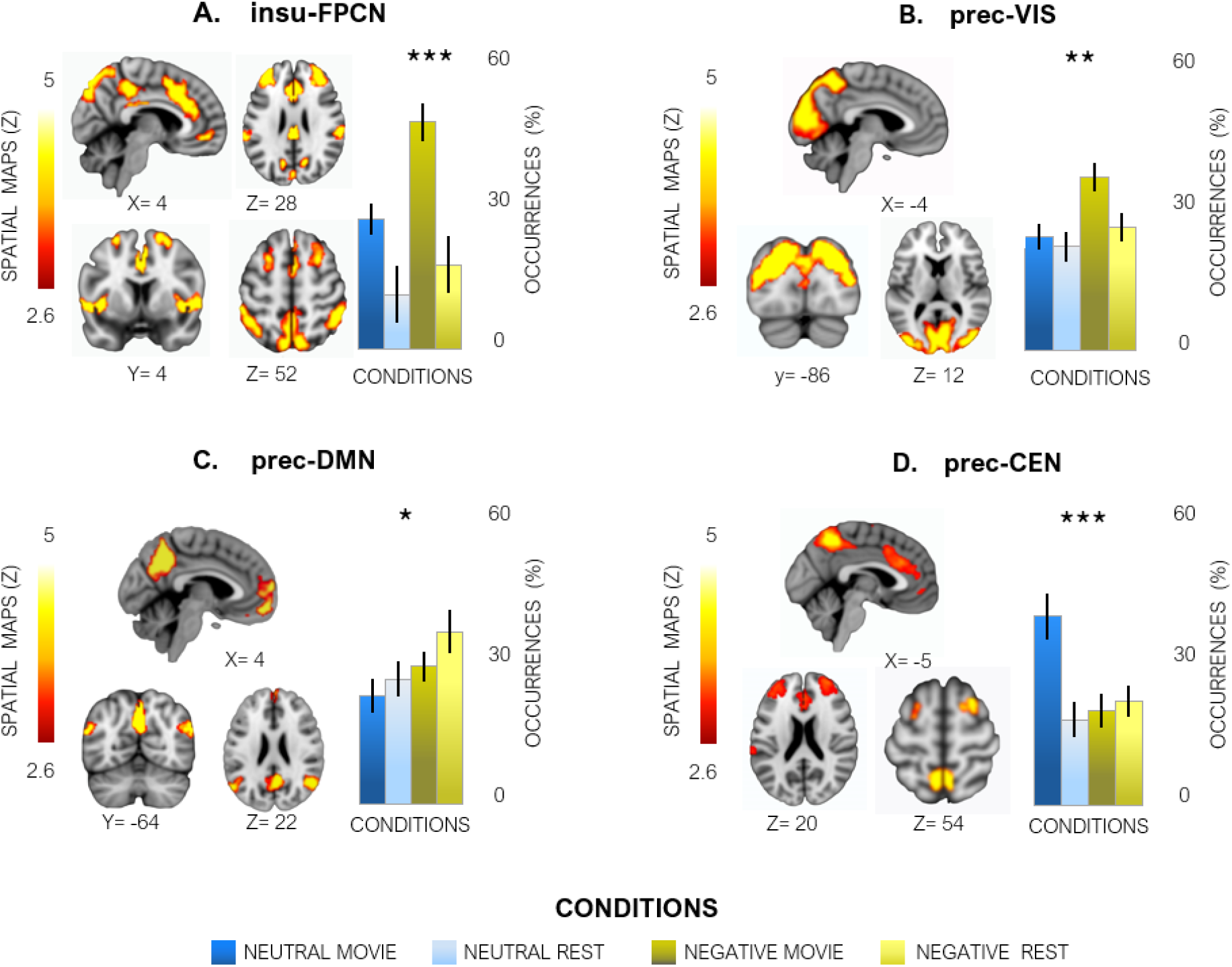
Spatiotemporal CAPs modulated by affective context. Functional brain networks were identified using co-activation pattern analysis (CAP) based on time-dependent coupling with the insula [A. insu-FPCN (frontoparietal control network)] or with the precuneus [B. prec-VIS(visual network), C. prec-DMN (default mode network), D. prec-CEN (central executive network)]. Only these four CAPs showed a signifcant modulation of their occurrence rate as a function of experimental condition (see suppl. Fig. S2). The brain topography maps of each CAP is illustrated with a threshold of z > 2.58 [equivalent to p<.01 (Liu et al., 2018)]. Further anatomical information on peak regions is provided in Table s1. Temporal occurrence rates are plotted to show their expression (%) across experimental conditions. Statistical assesments of CAPs variance and interactions are reported in Table 1., and Fig. S2. P_FDR_ adjustment for multiple comparisons (* p<.05; ** p<.01; ***p<.001).

Among networks co-activated with the insula, insuCAP5 was the only one showing a significant modulation by affective valence. It comprised a large set of frontoparietal regions associated with cognitive control and salience detection (Cole et al., 2014; Dixon et al., 2018a), and, therefore, considered to overlap with the FPCN reported in other studies (Figure 2A). This CAP exhibited an interaction between “affective context” and “stimulation period” [gLMM interaction: F(54)= 4.5; P .03], indicating more frequent occurrences of this network during negative movies compared to other conditions, in addition to main effects of both period [gLMM main effect “period”: F(54)= 46.6; P .<0001. β_movie_= 19.30, 95% CI (16.32; 22.30)], and context [gLMM main effect “context”: F(54)= 15.8; P .0002. β_negative_= 16.83, 95% CI (13.86; 19.80). Remarkably, 48% of the occurrences of this insuFPCN CAP across all the conditions were seen in the “negative movie” condition [in contrast to 24% during “neutral movie” (see post-hoc results in table 1)].

The precuneus-based precCAP1 overlapped with a classic “visual network” centered on occipital and posterior parietal areas (precVIS, Figure 2B) (Göttlich et al., 2017; Kang et al., 2016; Liu et al., 2018a; O’Connor et al., 2002; Riedel et al., 2018; Saarimäki et al., 2018; Sripada et al., 2014). It showed a distinctive increase of occurrence rates during the “negative movie” condition, with main effects of both “affective context” [gLMM main effect “context”: F(54) = 7.9; P =.006. β_negative_= 18.40, 95% CI (15.94; 20.90)] and “stimulation period” [gLMM main effect “period”: F(54)= 4.4; P =.03. β_movie_= 17.79, 95% CI (15.33; 20.30)], reflecting generally higher occurrences during movies than rest and during negative than neutral conditions, respectively. However, there was no significant “period” x “context” interaction [gLMM interaction: F(54) = 2.1; P=.15] despite its predominance during the negative movies, in which the precVIS exhibited 35% of total occurrences across conditions [vs. 21% during neutral movies (see post-hoc results in table 1)].

Two further emotion-sensitive CAPs were also derived from the precuneus-based analysis. Most notably, the precCAP3 showed a very similar spatial configuration to DMN (Christoff et al., 2016), encompassing medial prefrontal, posterior cingulate, and inferior parietal cortices (Figure 2C). The factorial analysis indicated a main effect of “affective context” [gLMM main effect “context”: F(54)= 4.9; P= .03; β_negative_= 16.7, 95% CI (13.85; 19.60)], but there was no significant effect of stimulation period [gLMM main effect “period”: F(54) = 2.2; P =.13; β_rest_= 16.74, 95% CI (13.18; 19.01)], nor any “context” x “period” interaction [gLMM interaction: F(54)= 0.27; P =.60]. However, this precDMN CAP dominated in the “negative rest” condition”, which represented 32% of its occurrences across conditions [vs. 22% during neutral rest (table 1)]. In contrast, the precCAP5 involved dorso-lateral fronto-parietal areas (Figure 2D) commonly associated with the central executive network (CEN) (Goldman-Rakic, 1995; Menon and Uddin, 2010). This network also showed a significant interaction between “affective context” and “stimulation period” [gLMM interaction: F(54)= 10.2; P= .002], elicited by a combination of increased occurrences in the “neutral” conditions [gLMM main effect “context”: F(54)= 4.5; P=.03. β_neutral_= 15.58, 95% CI (12.45; 18.70)] and in the “movie” conditions [gLMM main effect “period”: F(54) = 7.0; P=.01; βmov_i_e= 16.11, 95% CI (12.98; 19.20)]. Thus, unlike the precDMN_CAP3_, the precCEN_CAP5_ showed the highest number of occurrences during the “neutral movie” condition [39% of its total occurrence vs. 21% during negative movies (see statistical contrasts in table 1)].

Altogether, these data converge with previous studies to indicate that emotional events produce distinctive and coordinated effects on specific IFNs, especially the DMN and higher regulatory systems associated with FPCN and CEN, with significant carry-over effects lingering beyond emotional events themselves and extending into subsequent rest periods.

### Relationship of brain CAPs with behavioral measures of affect

To probe for the behavioral significance of changes observed in brain networks across conditions, we examined whether the individual occurrences of each relevant CAP could predict subjective affective rating scores (from PANAS) that were obtained at the end of scanning runs in each emotional context. Results from our GEE analysis showed that more negative affect (NA) scores post-scanning were associated with higher occurrences of insuFPCN CAP during the “negative movie” period (β = .24, 95% CI (0.15;0.33); P_FDR_ < 01; Figure 3B. Up-right), and inversely to lower occurrences of insuFPCN during the “negative rest” period (β = −.32, 95% CI (−0.30; 0.13); P_FDR_ < 05, Figure 3B. Bottom-right). On the other hand, occurrences of the precDMN CAP during the same “negative rest” condition also showed a positive correlation with the post-scanning NA scores (β = .20, 95% CI (0.20; 0.46), P_FDR_< 01). Figure 3B. Bottom-right). Thus, higher NA scores following exposure to the “negative context” were linked not only to greater occurrences of insuFPCN during negative movies, but also to lesser occurrences of insuFPCN accompanied with greater occurrences of precDMN during the subsequent “negative rest” period. By contrast, positive affect (PA) scores were not related to the expression of any of these networks (see Figure 3B. Left).

**Figure 3.**
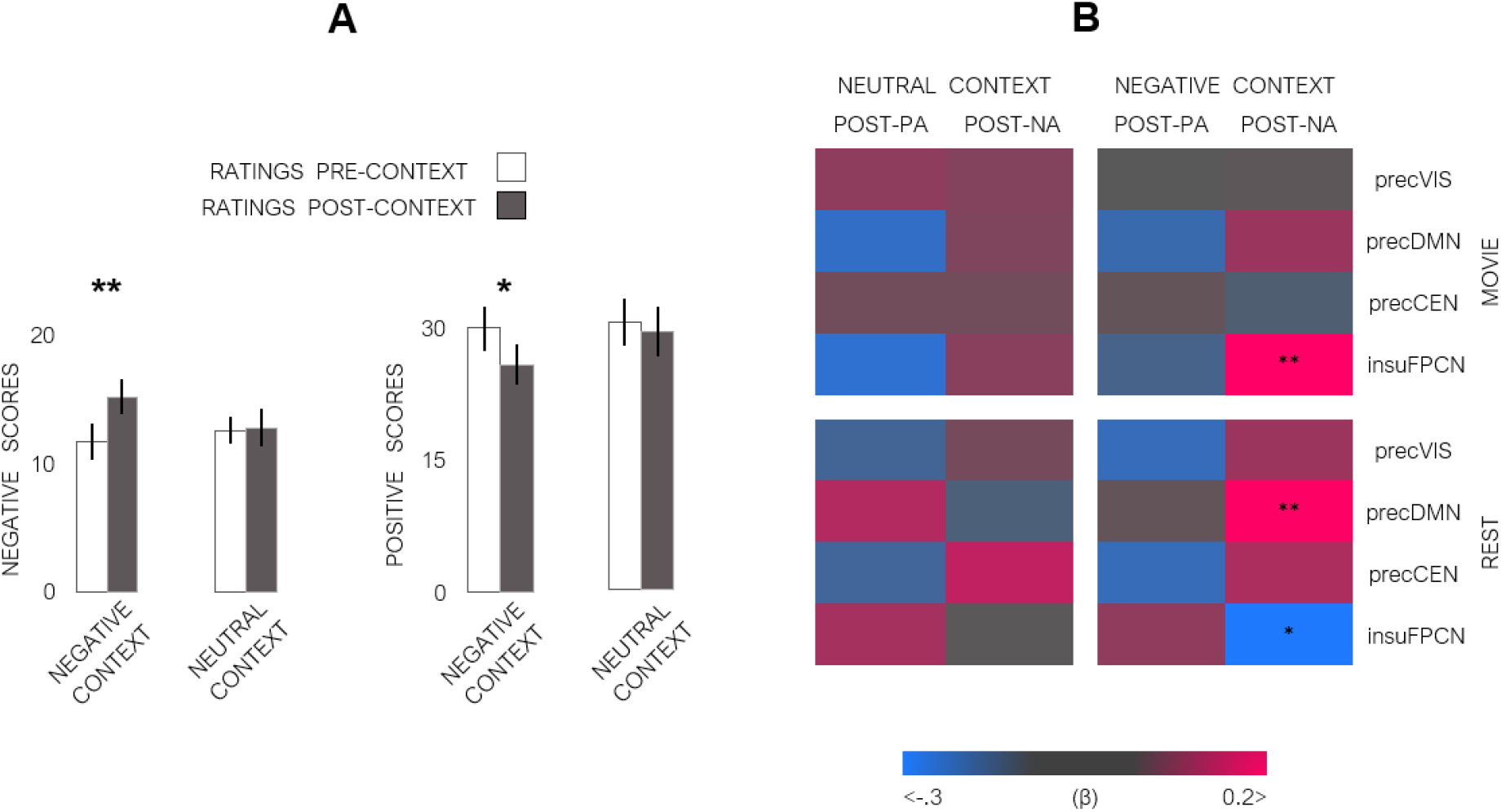
PANAS scores and their associations with CAPs across conditions. **A)** Mean scores on PANAS self-reports obtained before (pre) and after (post) each of the experimental contexts (neutral and negative). **B)** Beta (β) values of GEE-based associations between affective scores [positive (PA) and negative (NA)] and occurrence rates of the CAPs of interest. InsuFPCN occurrences observed during the “negative movie”, as well as insuFPCN and precDMN occurrences during the “negative rest” conditions were significantly correlated with the negative affect scores (NA) measured at the end (post) of the negative context. P_FDR_ adjustment for multiple comparisons (* p<.05; ** p<.01).

### Functional interactions within CAPs

Finally, we turned to the key question of our study. Namely, whether the reciprocal relationships between brain networks *during* and *after* movie episodes were modified by their affective valence (i.e., neutral or negative). To do so, we first computed movie-rest correlations in individual occurrences for each CAP of interest (precVIS, precDMN, precCEN, and insuFPCN) across all participants, and then examined how these relationships differed between the two affective contexts [within-CAP transitions; e.g., r(CAP(A)_neutral movie_; CAP(A)_neutral rest_) vs. r(CAP(A)_negative movie_; CAP(A)_negative rest_)]. Remarkably, we observed a consistent anticorrelation between the expression of a given CAP during movie periods and its expression during subsequent rest, for both affective contexts and for the precVIS, precDMN, and insuFPCN CAPs, but not for precCEN (see Figure 5A). Although these anticorrelations were numerically higher in the negative context (r values −.46 to −.67) than the neutral context (−.33 to −.39), a direct comparison of anticorrelation magnitudes (see methods) showed no significant difference for any of the four CAPs.

However, the within-CAP correlations for transitions from movie to rest conditions were significantly modulated by affective context in comparison to control conditions when the movies and rest did not follow in direct succession [e.g., r(CAP(A)_neutral movie_; CAP(A)_negative rest_], but selectively for the precDMN (Figure 4) and not for the three other networks. Thus, for the precDMN CAP, the anticorrelation pattern observed during the post “negative movie” transition to the subsequent “negative rest” condition across all participants [r(19)=−.67; P= 0009; P_FDR_=.01] was significantly higher relative to the anticorrelation between non-successive “negative movie” and “neutral rest” periods [r(19)=−.19; P=.38 P_FDR_= .38. P=.02, CI=(−1.16; −.03) for the difference], or relative to the anticorrelation observed between non-successive neutral movie and negative rest periods [r(19)=−.05; P_FDR_<1. P=.04, CI=(−1.14 −.01) for the difference]. None of the other CAPs showed any difference in these comparisons (Table S2).

**Figure 4.**
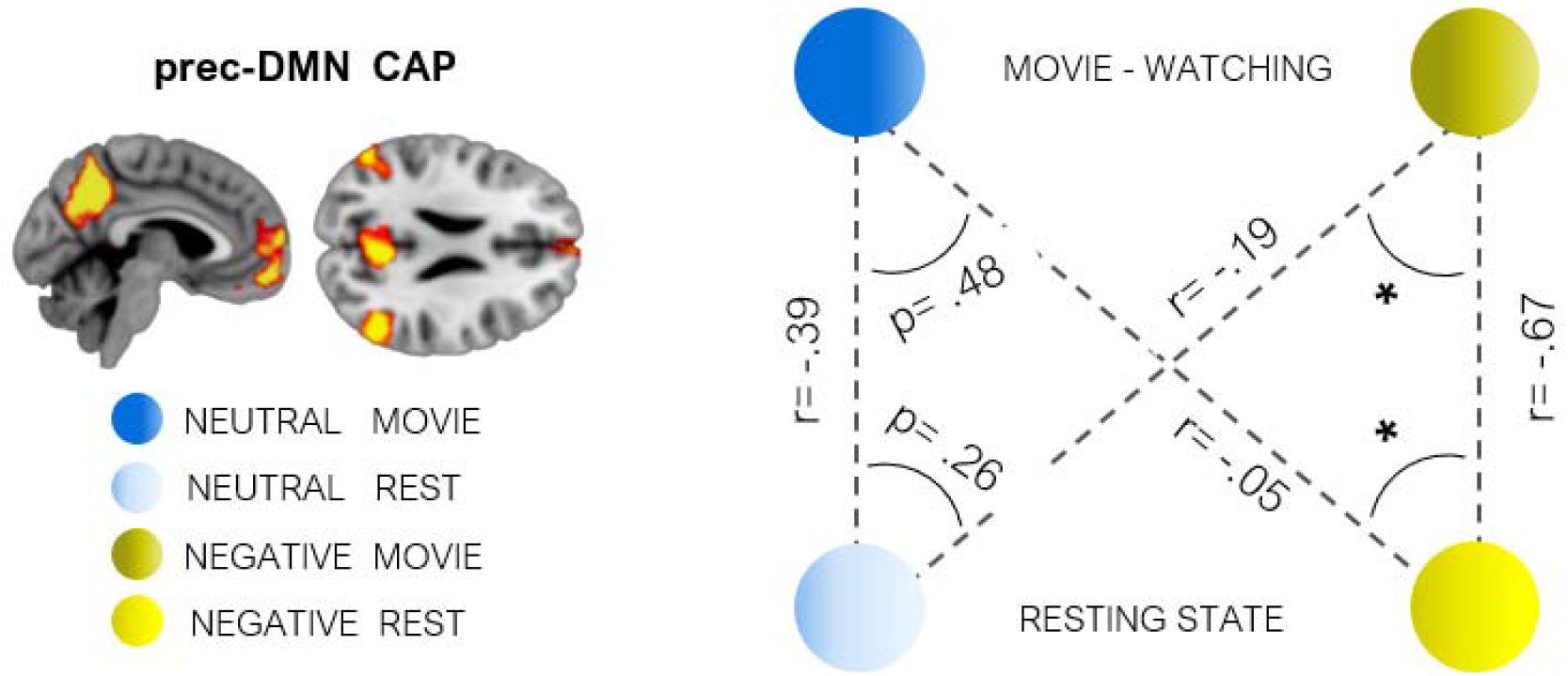
Within-DMN CAP interactions across. Comparisons of the movie-rest anticorrelation for the DMN (temporal occurrences per condition), illustrating significant differences for transitions between successive movie-rest periods in the negative context (yellow colors), relative to control comparisons with conditions from the “neutral” context (blue colors). The other three CAPs of interest (prec-VIS, prec-CEN, insu-FPCN) showed no such effect of negative emotion on the same transitions. * p<.05. Dependency of correlations across conditions was considered when comparisons included overlapping variables (Zou, 2007).

Altogether, these data highlight robust antagonistic relationships between the expression of particular functional networks during active movie watching and their subsequent expression during rest, regardless of affective state (except for the precCEN CAP), but with a distinctive impact of negative emotion on these transitions for the precDMN CAP (Figure 4).

### Functional interactions between CAPs

Lastly, we examined reciprocal interactions among networks by determining whether the occurrence rate of a given CAP of interest during movie watching could predict the occurrence of *other* CAPs during the following rest period, and tested whether this relationship varied in a context-dependent manner for negative compared to neutral conditions (between-CAP analysis; e.g., r(CAP(A)_negative movie_; CAP(B)_negative rest_) vs. r(CAP(A)_neutral movie_; CAP(B)_neutral rest_)]. Results from this analysis (Figure 5B) revealed a positive association between the insuFPCN in the moviewatching period and precCEN in subsequent resting period in the “neutral” context [r(19)= .54; P=.02; P_FDR_=.04], which was significantly different [P=.02; CI=(.02; 1.23). Figure 5B.] from the suppression of such interaction in the “negative” context [r(19)=.-19; P=.44; P_FDR_=.44]. Conversely, the insuFPCN during movie watching exhibited a positive correlation with the precDMN during subsequent rest exclusively in the “negative” context [r(19)=.28; P=.23; P_FDR_=.23], absent in the “neutral” context [r(19)=−.34; P=14; P_FDR_=.14], with a significant difference between the two contexts [P =.04; CI=(−1.17; −0.10). Figure 5B]. On the other hand, the precDMN during movie watching was strongly anticorrelated with the precCEN during subsequent rest exclusively in the “negative” context [r(19)=.-61; P=.003 P_FDR_=.005], a significant difference [P=.02; CI=(0.02; 1.20)] compared to the absence of such correlation in the neutral context [r(19)=−.11; P=.64; P_FDR_=.64 (Figure 5B)]. Other relationships between pairs of networks were not significant and not modulated by affective context.

**Figure 5.**
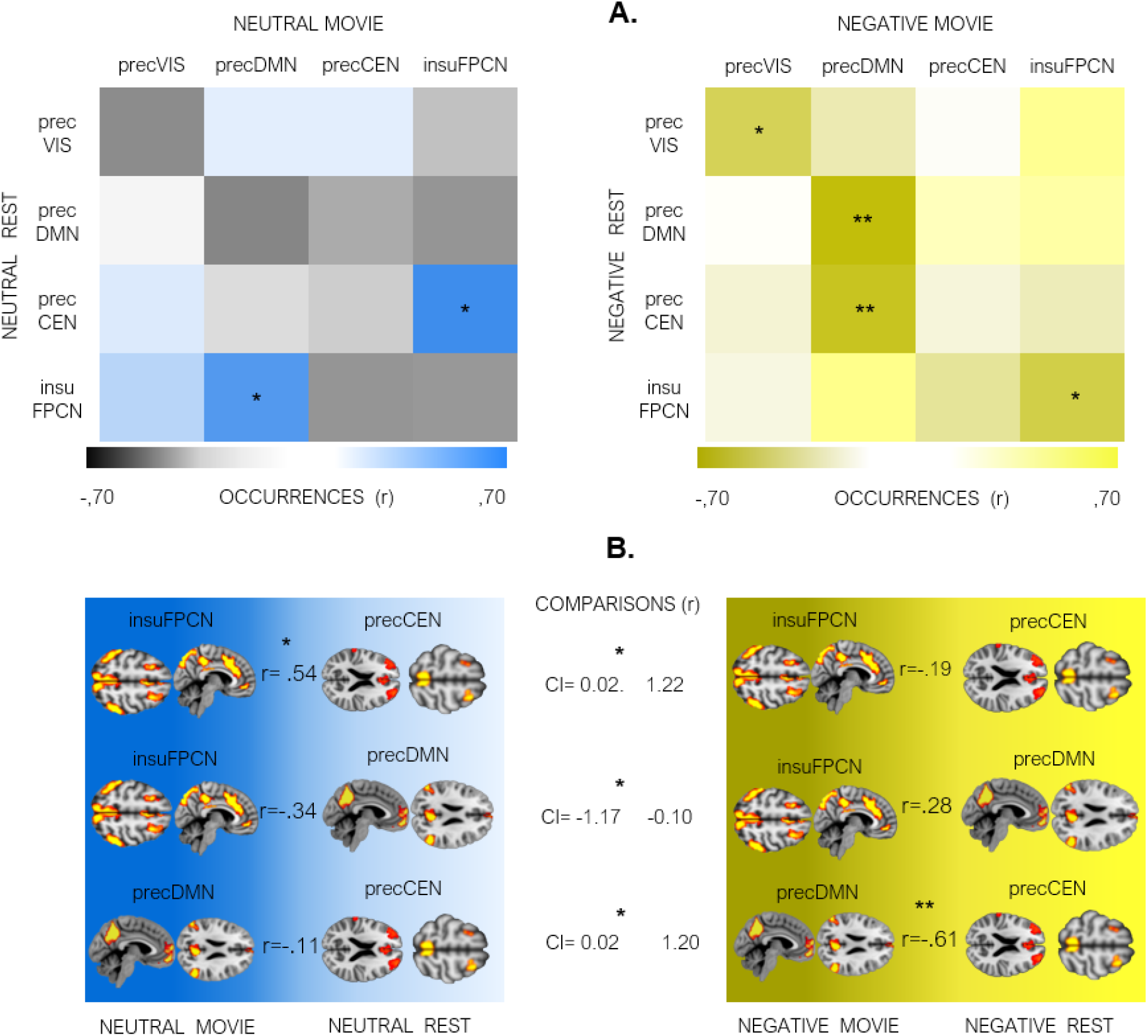
Between-CAPs interactions across time. **A.** Pairwise correlations in occurrences between CAPs of interest observed during movies and rest periods, respectively, computed for the “neutral” (left) and the “negative” context (right), P_FDR_ adjustment for multiple comparisons (* p<0.05; ** p<0.01). **B.** Comparisons of between-CAPs correlations that showed significant differences between the two emotion contexts. Dependency of non-overlapping variables were considered following (Zou, 2007).

## Discussion

Our study identified a set of distributed brain networks with dynamically fluctuating co-activation patterns (Bolton et al., 2020; Liu et al., 2018b; Preti et al., 2017) that were modulated by negative emotion, with differential expression (i.e., occurrences) and interactions (i.e., correlations) observed both during emotional stimulation itself (sad movies) and during its aftermath (subsequent rest). Specifically, we uncovered four networks whose temporal profile differed as a function of emotional conditions, overlapping with the DMN, CEN, FPCN, and VIS networks reported in other studies. These results show, first, that negative emotional events engage a select set of brain-wide systems, with anatomically and temporally specific connectivity arising not only during emotional events, but also extending during subsequent rest (Figure 2). Second, the occurrences of some of these CAPs is directly related to subjective emotional state, with more negative affect (NA) reported by participants who show more frequent occurrences of the insuFPCN and the precDMN during and after negative movies, respectively (Figure 3). Third, we observed specific temporal relationships “within” and “between” these networks, such that their expression rate during movie periods predicted their expression in subsequent rest. Critically, these functional dynamics among networks differed significantly in the negative affective compared to the neutral conditions (Figures. 4-5). Together, these data provide new evidence on how brain-wide systems are dynamically and interactively recruited by emotions and their regulation, with robust carry-over effects beyond emotional events themselves. These neural results add to behavioral evidence of “emotional inertia” (Kuppens et al., 2010; Kuppens and Verduyn, 2017) and suggest that emotion regulation processes may extend over protracted periods of time following emotion elicitation. More generally, our study provides new insights on affective brain dynamics and uncovers useful neural markers for adaptive regulation mechanisms that mediate a restoration of homeostatic balance after stressful events, possibly altered in psychopathological conditions such as depression, anxiety, or PTSD.

### Emotion-responsive CAPs with different temporal profiles and different functional roles

Among networks differentially modulated by negative emotion, the precVIS CAP comprised visual areas (Orban et al., 2004) mainly located in dorsal extrastriate cortex and mostly engaged during the “negative movie” periods. This accords with abundant evidence for enhanced activity of the visual system in response to affective stimulation (Kragel et al., 2019; Vuilleumier et al., 2004), presumably reflecting modulatory top-down signals from emotion-responsive limbic areas (McAlonan et al., 2008; Vuilleumier and Driver, 2007). Nevertheless, precVIS activity showed no direct interaction with the expression of other CAPs of interest, consistent with these sensory areas having no direct link with emotional experience per se. On the other hand, the insuFPCN CAP also predominated during emotion elicitation periods (negative movie), but its occurrences correlated with negative affect scores reported after scanning. Anatomically, insuFPCN comprised components of the “salience” and “dorsal attention” systems, typically engaged in the appraisal of arousing information (Sander et al., 2018). This pattern fits well with theoretical accounts (Feldman B. and Satpute, 2013; Scherer and Moors, 2019) according to which emotion involves an integration of processes associated with selective attention (Touroutoglou et al., 2012), interoception (Critchley and Garfinkel, 2017), and detection of behavioral relevance (Roy et al., 2012). Higher insuFPCN expression during the negative movie indicates this network is an important component of neural processes underlying emotion elicitation (Meaux and Vuilleumier, 2015; Sander et al., 2018).

In contrast, the precDMN CAP showed the greatest number of occurrences during rest periods following negative movies, and its presence also correlated with negative affect scores. These results add to recent findings of similar aftermaths of emotionally arousing stimuli during resting state, e.g., following fearful or joyful movies (Eryilmaz et al., 2011) as well as reward or punishment outcomes (Eryilmaz et al., 2014; Pichon et al., 2015). Emotional carry-over effects in brain connectivity may also underpin changes in affective and cognitive responses to future events (Qiao-Tasserit et al., 2017), and thus contribute to adaptive behavioral functions of emotions. Enhanced precDMN activity at rest has previously been associated with self-referential mental activity and introspection (Christoff et al., 2016; Fox et al., 2018), autobiographical memory (Engen et al., 2017), as well as mood disorders (Liemburg et al., 2012; Sheline et al., 2009; Song et al., 2016). PrecDMN upregulation is often regarded as a biomarker for depressive rumination (Hamilton et al., 2015; Sambataro et al., 2014; Whitfield-gabrieli and Ford, 2012), possibly resulting from emotion dysregulation (Aubry et al., 2016). Moreover, both spontaneous emotion regulation strategies (Abler et al., 2010; Joormann and Gotlib, 2010) and voluntary cognitive reappraisal (Ertl et al., 2013; Uchida et al., 2015) enhance DMN connectivity, suggesting a link with functional recovery mechanisms acting to downregulate negative emotions as observed in the post sad movie rest periods here.

Additionally, in our study, precDMN was the only CAP exhibiting a significant impact of the negative affective context on functional transitions between its expression rate during rest and its expression during the just preceding movie (within-CAP dynamics). Although a consistent anticorrelation of CAP occurrences between movie and rest periods (i.e., more frequent occurrences during movies predicting less frequent occurrences during rest) was observed for all emotion-responsive networks (except precCEN), only the precDMN showed a statistically significant amplification of this antagonistic relationship for negative rest periods that directly followed negative movie periods. In other words, the lower the expression of precDMN during the sad movie, the higher its expression during subsequent aftermath at rest (see Figure 4). This strong and valence-dependent shift in dFC provides new evidence for a key role of the DMN in homeostatic adjustment after acute stressors (Hermans et al., 2014), possibly through coordinated interactions with other brain networks (see below). Anti-correlated activity among networks may represent a general and biologically meaningful process in functional brain dynamics across various conditions (Li et al., 2021). Nevertheless, it remains to fully determine the factors accounting for such “push-pull” DMN anticorrelations in occurrences between emotional episodes and subsequent recovery periods at rest. Importantly, the absence of such behavior in movie-rest transitions for the precCEN indicates that this functional shift in connectivity state is not caused by non-specific rebound or hemodynamic effects but appear both network-specific and context-sensitive.

Finally, the precCEN showed a distinctive profile in occurrences, with predominant expression during neutral movies relative to other conditions. This CAP comprised dorsal ACC and dorsolateral prefrontal areas associated with effortful executive control (Cole et al., 2013), presumably activated by more abstract cognitive demands to process the content of neutral movies (unlike more natural absorption by emotional scenes). Nevertheless, the precCEN exhibited significant changes in its interaction with other CAPs as a function of affective context, as demonstrated by our analysis of between-CAPs relationship and discussed below.

### Emotion-related modulation of functional interactions between CAPs

A key finding of our study is that higher or lower expression of certain CAPs during the movie period was associated with different occurrences of *<u>other</u>* CAPs during the subsequent rest period (between-CAPs temporal dynamics), and these relationships among networks were significantly modified by the emotional context. Most critically, negative emotion produced a significant shift in the functional impact of insuFPCN activity on subsequent precDMN and precCEN at rest, such that a strong relationship of insuFPCN with precCEN in the neutral context was suppressed in the negative context, and replaced by a positive relationship with precDMN instead (Figure 5B). Thus, higher occurrences of insuFPCN <u>during</u> sad movies (which correlated with more negative affect) predicted greater presence of the precDMN at rest <u>following</u> these movies (which also correlated with negative affect). This suggests that higher affective salience encoded by insuFPCN might enhance subsequent self-regulatory processes and ruminative thoughts subserved by precDMN (Andrews-Hanna et al., 2014). We note that the magnitude of correlations between insuFPCN and subsequent precDMN activity was numerically modest in both affective contexts but shifted from a negative relationship in the neutral context (r= −.34) to a positive relationship in the negative context (r=.28), a highly significant difference between these two conditions.

In parallel, precCEN expression at rest also shifted from a strong positive correlation with insuFPCN occurrences during the preceding movie in the neutral context, to a strong negative correlation with precDMN occurrences in the negative context, both representing highly significant changes in correlation magnitude and/or direction. These opposing effects on precCEN may accord with behavioral evidence for impaired executive control abilities after acute stress conditions (Chand et al., 2017), which could be mediated at least partly by inhibitory interactions from precDMN in the post-negative context. Conversely, the positive relationship between insuFPCN and subsequent precCEN at rest in the neutral context suggests more synergic interactions between these two networks, commonly implicated in externally oriented cognition (Corbetta and Shulman, 2002; Spreng et al., 2016). Interestingly, such modulation of precCEN expression by preceding movie valence, despite low occurrences of precCEN at rest overall (Figure 2D), underscores that a high occurrence rate is not a prerequisite for measuring significant changes in CAP dynamics, and further shows that emotion may impact on the functional integration of brain networks rather than just their global activation level (Leitão et al., 2020).

In sum, our findings show not only distinctive patterns of expression of insuFPCN and precDMN CAPs during negative emotion episodes and post-emotion rest periods, respectively, but also significant functional relationships between these networks during the transition from emotion episodes to subsequent rest. Together, these data point to complementary roles of FPCN in the integration of emotion elicitation processes and of DMN in self-regulation mechanisms contributing to affective homeostasis.

### Outstanding issues and further directions

Although we found robust differences in the expression and correlation of relevant CAPs in our study, and carefully counterbalanced conditions between participants, we cannot fully rule out that other effects related to movie contents or time contributed to changes in network activity and their functional relationships as observed here. Further research using other naturalistic designs and evoking different emotions (e.g., frustration, joy, etc.) will also be valuable to confirm and extend our results. In addition, our gender selection was limited to female participants, based on pilot data showing higher and more consistent emotion ratings than males, and previous studies showing more intense emotion experiences in females (Vrtička et al., 2012). It will be important to replicate and extend the current findings to male populations, because emotion dysregulation and resilience to stress may show important gender differences.

### Conclusions

Using a recent dFC analysis methodology (CAPs), we shed new light on the spatio-temporal organization of brain networks underlying the dynamics of negative affective experience and subsequent return to rest. Our data unveil specific transitions in the connectivity and reciprocal relationship of brain-wide networks involved in visual perception, attention to salient stimuli, executive control, and introspective self-monitoring processes during both emotional episodes (movies) and their aftermath (subsequent rest). Among these networks, the insuFPCN and precDMN CAPs were found to play a pivotal role in the temporal dynamics of emotional experience, from elicitation to subsequent recovery, entertaining reciprocal relationships not only with each other but also with the precCEN CAP. Crucially, individual occurrences of these CAPs during movies and rest were directly linked to the subjective affective state reported by participants. These findings provide novel insights on brain mechanisms underlying emotion experience and regulation (Dixon et al., 2018b; Pan et al., 2018; Sripada et al., 2014), resilience to stress (Young et al., 2017), and perseverative ruminative thinking in negative mood states (Eryilmaz et al., 2011; Garfinkel et al., 2016; Harrison et al., 2008). In turn, they may also offer valuable biomarkers for understanding and assessing the neural basis of clinical psychopathology conditions. Nevertheless, because our sample was restricted to female participants, our results constitute a preliminary exploration of brain networks dynamics in response to sad emotions, and future research in larger, independent samples is needed to generalize our findings.

## Acknowledgments

This work was supported by the Swiss excellence Scholarship program, the Schmidheiny foundation, and the Colombian Science Ministry (JG), as well as by a Sinergia Grant no. 180319 from the Swiss National Science Foundation (SNF), by the Swiss Center of Affective Sciences financed by UNIGE and SNF (Grant no. 51NF40_104897), and by the Société Académique de Genève (SACAD). This study was conducted on the imaging platform at the Brain and Behavior Lab (BBL) and benefited from support of the BBL technical staff. Special thanks to dr. Frédéric Grouiller for his useful comments on the imaging processing analysis. Imaging analysis was carried out at University of Geneva on the high-performance computing (HPC) BAOBAB server.

